# TopoTome: Topology-informed unsupervised segmentation and analysis of 3D images

**DOI:** 10.1101/2025.09.26.678737

**Authors:** Samuel Bourgeat, Fani Derveni, Victoire Gamblin, Ece Naz Bilgiç, Feyza Nur Arslan, Andrew Oates, Can Aztekin, Fides Zenk, Pedro M. Reis, Ana Marija Jakšić

## Abstract

Biological systems are three-dimensional and complex. Today, images of biological structures can be acquired using different imaging technologies and at increasing resolutions. However, identifying relevant structural features in three-dimensional (3D) images remains a significant challenge. 3D image segmentation is usually performed using deep-learning segmentation models. Such models are trained on manually annotated and segmented, dataset-specific images. Consequently, they rarely generalize across datasets. Here, we overcome these limitations with TopoTome, an unsupervised 3D image segmentation and analysis algorithm. TopoTome is based on topological data analysis, and conceptually distinct from standard clustering and deep learning image analysis models. It encodes the 3D image directly in topological space. Then, it performs unsupervised clustering to detect and segment features represented by salient and spatially coherent voxel intensity gradients. We demonstrate on simple, complex, synthetic and real-world 3D image data that it outperforms all 3D clustering algorithms. Its segmentation ranks with or outperforms best-in-class deep learning 3D image segmentation software. Beyond image segmentation, it provides streamlined topological data analysis of 3D images, advancing 3D image analysis from conventional mesh volumetry to structural topology. Owing to its conceptually different topological data analysis core, TopoTome does not need prior information and tuning to generalize across different imaging modalities, including fluorescence microscopy and X-ray computed tomography. We show it also readily generalizes across biological subjects, such as different species, organs and cells. TopoTome is thus one of the most versatile and accurate unsupervised 3D image segmentation algorithms.

## Introduction

Biological systems are composed of intricately organized three-dimensional structures that span multiple organizational scales, from molecules, cells, and tissues to organs. Recent advances in imaging technologies, including light-sheet microscopy^1–3^ and X-ray computed tomography^4,5^ have made it possible to capture these structures in rich 3D detail. The analytical bottleneck has since shifted from data acquisition to data analysis and interpretation. Processing and analysing complex 3D image data remains a formidable challenge in bio-imaging.

Artificial intelligence (AI) and, more precisely, deep learning (DL) methods have enabled substantial progress in that respect, particularly in accelerating segmentation and feature extraction from well-structured datasets. However, these approaches are still heavily reliant on vast amounts of data and ground truth annotations generated through human input^6–9^. Such annotations are labour-intensive, subjective, and difficult to obtain consistently for large, heterogeneous, or highly complex data. This is especially difficult for 3D images^10,11^.

The dependency on manual data curation introduces a critical limitation: models trained on sparse or human-preprocessed datasets are dataset-specific and fail to generalize to image data with different statistics. The limitation stems from variable levels of image complexity, data noise, or variable imaging parameters and modalities. Further, despite being termed “the ground truth”, manual segmentations are inherently influenced by human bias and our perceptual constraints. This is particularly problematic when dealing with dense or low-resolution volumetric data where object boundaries appear ambiguous to the human eye or span multiple stacked 2D images^12^. Unsupervised, deterministic algorithms such as intensity-based thresholding or clustering methods are commonly employed to assist humans in deciding on the ground truth. For example, binarising images with tuned thresholding helps with finding coarse feature boundaries in images that lack obvious, sharp intensity gradients. Unsurprisingly, such unsupervised algorithms often perform as well as even the best trained 3D segmentation algorithms when applied to non-challenging, “clean” datasets^7^. Yet automated thresholding falls short when applied to morphologically complex, non-homogeneous biological architectures, or simple but low-resolution, noisy datasets.

Consequently, current standard image analysis pipelines first use unsupervised or semi-supervised image preprocessing, including human ground truth definition, neural network design, model training and optimization (**Figure 1A**). After the training, final automated segmentation is performed on the full dataset, and is usually followed by extensive manual post-processing to correct segmentation errors^10,12^. Even such elaborate workflows struggle to deliver robust, scalable, and reproducible segmentations if the 3D datasets are sparse, heterogeneous, or complex - a rule, rather than an exception, in the realm of biology. These limitations underscore the need for a new methodological framework that can go beyond human intuition and conventional heuristics. Such a method would ideally enable an entirely algorithmic analysis of biological 3D structures.

**Figure 1.**
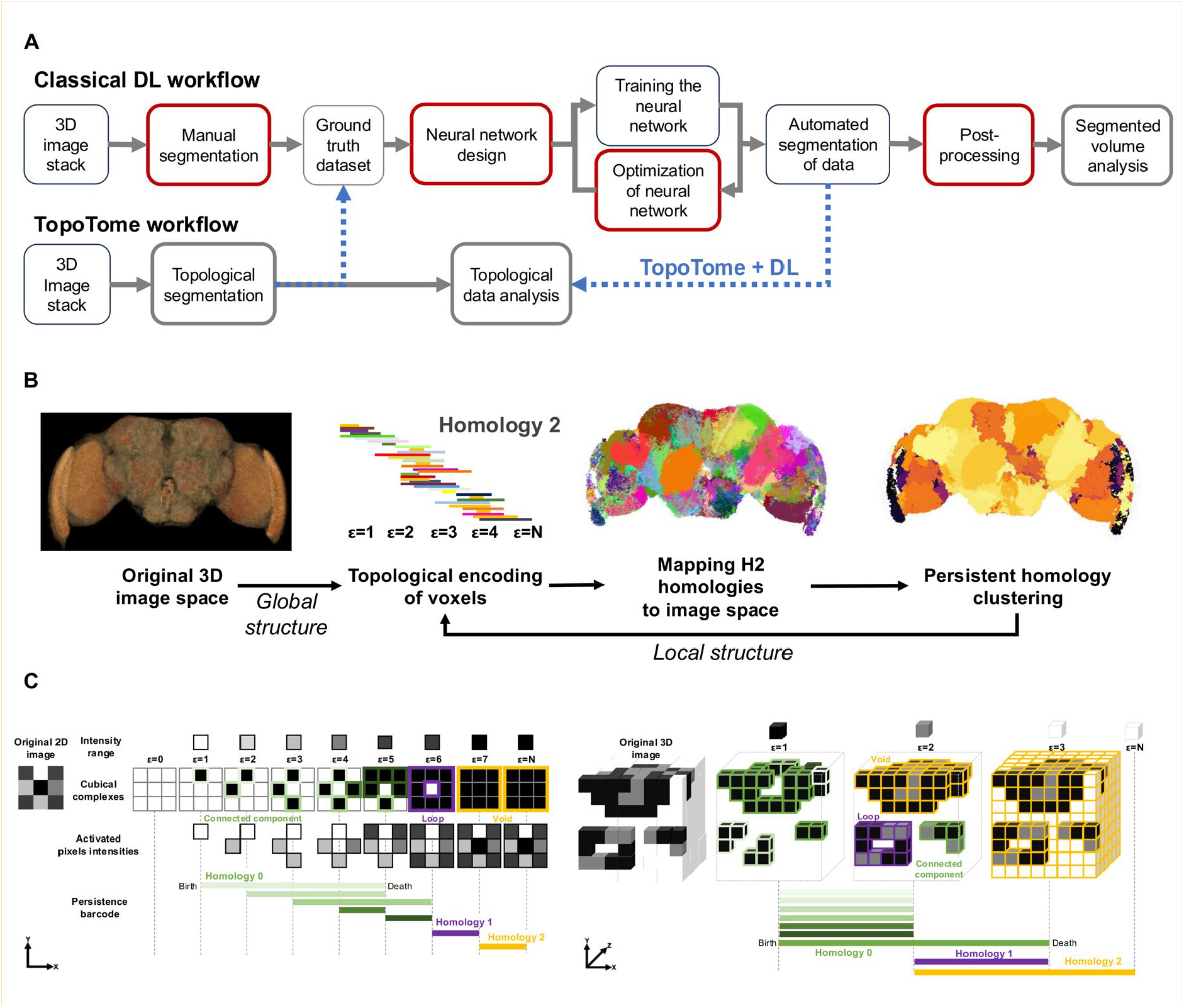
TopoTome: topological data analysis and unsupervised morphometrics of brain 3D tomograms. **A)** Deep learning segmentation workflow vs. TopoTome workflow. Red boxes indicate steps that require lengthy manual intervention of the experimenter. Synergistic workflow between DL and TopoTome in which TopoTome segmentation is used for determining ground truth for DL model training is proposed in blue. **B)** Core concept underlying TopoTome algorithm. **C)** Cubical complex filtration schematic on 2D and 3D greyscale images and the construction of H0 (green), H1 (purple) and H2 (yellow) persistent homologies (bars). Tracking the birth and death of homologies across filtration steps (*epsilon*) builds the persistence barcode. Here, epsilon denotes the maximum value of the considered voxel intensity range.

We address this need by developing TopoTome, a direct encoding of tomographic voxel data in topological space, and subsequent decoding of topological features back into image space (**Figure 1B**). Instead of relying on human annotations, TopoTome enables the identification of distinct volume features in 3D image data based solely on the topology of the distribution of voxel intensities in 3D space. It is readily applied to non-binarized structural 3D imaging data, and its output can be directly analyzed using topological data analysis (**Figure 1A**). It is not a deep learning algorithm, hence it does not require a ground truth dataset or model training. We show that TopoTome can be directly applied to segment non-trivial biological structures. This includes brain structures across scales of biological organization, including brain region segmentation across different species, organoid tissue segmentation, cell segmentation, and segmentation of non-biological 3D structures. Our findings establish the direct topological encoding of 3D images as a novel framework for unsupervised segmentation, but also as a novel concept for a quantitative topological analysis of complex 3D structures that moves beyond volumetry.

## Results

### Direct topological encoding of 3D images

Topological data analysis (TDA) is used to construct a topological representation of any shape or structure. It has been applied to various biological problems with promising results^13–19^. In most TDA use cases, the subject dataset is first translated into a point cloud dataset where the distance between points reflects some salient data property that we wish to analyse. To reduce the downstream computational load, large point clouds are typically subsampled and then transformed into a topological space using a complex filtration procedure (usually simplicial complex filtration). While this approach is computationally efficient, its main disadvantage comes from transforming data into distances before calculating its topology. This discards an amount of structural information in the data that may be crucial for identifying meaningful topological features. It also prevents direct mapping of topology back to the original dataset.

In TopoTome, we make use of the fact that 3D images are already grids of cube-shaped points - voxels. We can naturally represent them as cubes and directly encode them topologically using *cubical complex filtration* on their intensity (**Figure 1B, Figure 1C**)^20^. This way, rather than transforming the image data to a point cloud, we achieve a direct mapping between image space and topological space (**Figure 1B**). All downstream computation and image analysis is then applied to the topological representation of the image instead of the distances between points in the Euclidean space.

We use cubical complex filtration to identify independent and coherent 1D, 2D, and 3D shapes based on both local and global spatial distributions of voxel intensity values (**Figure 1C**, see Methods: Topological data analysis underlying TopoTome). This enables the use of topological data analysis of the entire image and its localized topologically relevant parts (**Figure 1A, 1B**). During the cubical complex filtration, we first consider an increasing voxel intensity range (the filtration step, *epsilon*) and analyze the spatial distribution of assemblies of voxels that fall within the range (**Figure 1C**). Assemblies of voxels will form distinct shapes across spatial dimensions, called *cubical complexes*. As the intensity range (*ε*) increases, we capture the topology of the evolving shape of these voxel assemblies. We do so by computing the persistence of cubical complexes across filtration steps. Their persistence is measured by their appearance (birth) and disappearance (death). As they are born and die, cubical complexes give rise to explicit topological features called *homologies*. Homologies are quantified across different topological dimensions as connected components (H0), loops (H1), or holes/voids (H2)^20^ (see details in Methods; **Figure 1C**). Together, the collection of these features forms the *persistence barcode*, a direct and analytically accessible encoding of the topology of the entire 3D image (**Figure 1C**).

The computation of a persistence barcode is deterministic and entirely data-driven, therefore reproducible. It can be quantitatively analysed using any standard algebraic topology toolkit (e.g.^14,21,22^). Importantly, persistence barcode construction is not constrained by structural diversity, complexity of the shape or image, or prior knowledge of the image statistics or imaged structure. This makes it especially well-suited to characterize the most intricate and unique biological structures. Direct topological encoding of the volumetric image underpins TopoTome algorithm’s core functions, including segmentation, quantification, and analysis of 3D features (**Figure 1A, Supplementary figure 1, Supplementary table 1**).

To map the persistent homology groups back on the original 3D data space, we use an algorithmic approximation of the calculation of local homologies. This feature is crucial for the biological interpretability of what the image topology represents and where it can be localized in the original image. When we use TopoTome to segment 3D features that enclose a volume, an explicit mapping procedure takes a pair of persistent H2 homologies from the persistence diagram and then maps them to the associated cluster of voxels in the original image (**Figure 1B**). This mapping is essentially an unsupervised segmentation task in which voxels from a 3D image are clustered based on their contribution to a topologically continuous, persistent feature. We then apply an additional level of standard Leiden clustering on the topological persistence clusters to identify local communities of topological clusters that represent a larger common structure (**Figure 1B, Supplementary figure 1**). The shape and topology of local communities can then be further analyzed, separately from the global topology of the image (**Figure 1B**).

Topological segmentation uses the continuity in local voxel intensity gradient in the context of the global distribution of voxel intensities. This mimics how humans decide where the structural boundaries of an object are, using local information as well as the global context of the entire image. However, in this case, the boundaries and the context are not perceived by eye, and the decision is not made probabilistically through human cognitive process or intuition. Rather, the global and local context is explicitly measured and quantified using the topology of the data, and the decision is made exclusively based on data structure. Moreover, the topology of the image is by definition a measurement of the invariant shape of the imaged object. This makes topological encoding useful not only for segmentation, but also for the downstream topological analysis of the shape of the imaged object.

### Benchmark against 3D clustering algorithms

We produced several synthetic 3D datasets to test the segmentation fidelity and runtime of TopoTome. We compared TopoTome to the most commonly used unsupervised clustering methods that can handle 3D image data, including K-means^23^, HDBSCAN^24^, Spectral^25^, and Agglomerative clustering^26^. We also compared it to the currently only other topology-informed clustering method, ToMATo^19^. We challenged the robustness of these methods and TopoTome by perturbing the original dataset with Gaussian, Salt and Pepper, Speckle, and Poisson noise. These perturbations represent common challenges in microscopy and medical imaging pipelines^27^. Compared to all the methods, TopoTome consistently produced high-fidelity 2D and 3D segmentation and was largely robust to all types of noise (**Figure 2, Supplementary figure 2A, 2B**).

**Figure 2.**
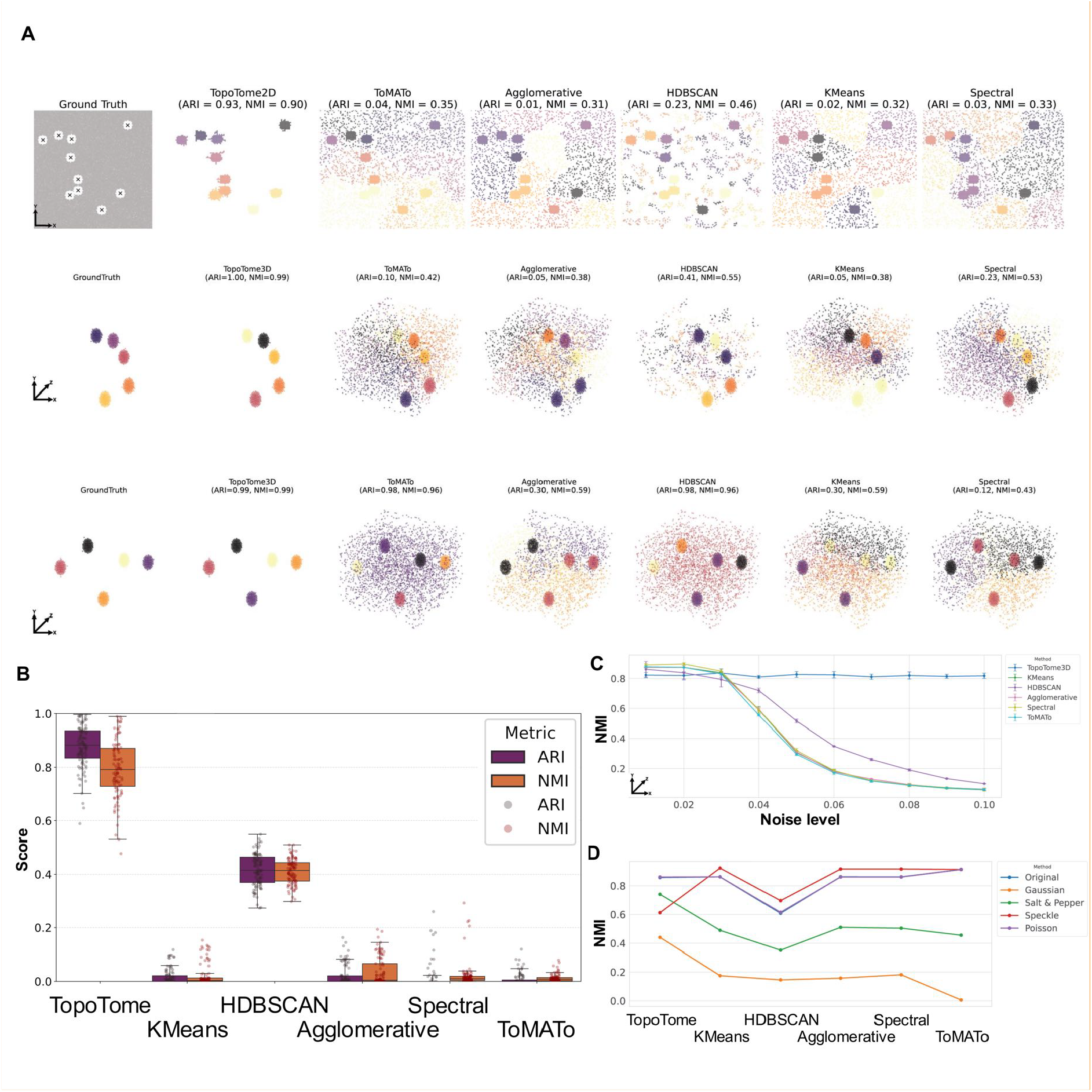
TopoTome benchmark against unsupervised clustering methods. **A)** Benchmarking TopoTome against clustering algorithms on 2D (top panel row) and 3D synthetic datasets using feature vectors based on spatial coordinates (middle panel row) or using feature vectors combining coordinates and voxel intensities (bottom panel row), all with high Gaussian noise (σ*s* = 0.1). **B)** Distribution of Adjusted Rand Index (ARI) and Normalised Mutual Information (NMI) performance scores over 100 runs for each algorithm on 2D images with moderate noise (σ = 0.05). TopoTome significantly outperforms all clustering algorithms. **C)** NMI values for each algorithm across 5 3D images per noise level (noise ranging from σ = 0.01 to σ = 0.1). **D)** NMI values across different noise distributions. TopoTome shows consistent robustness to noise, including Salt and Pepper and Gaussian noise.

For the Gaussian noise condition (σ = 0.1), TopoTome accurately recovered the ground-truth clusters with Adjusted Rand Index ARI = 0.93 and Normalized Mutual Information NMI = 0.90, while all other algorithms dropped well below 0.5 on both metrics (**Figure 2B**). Under moderate noise levels (σ = 0.05), repeated 100 times, TopoTome maintained robust clustering with ARI and NMI consistently in the 0.7-1.0

range (**Figure 2C, Supplementary Figure 2A**). In contrast, K-Means and Spectral Clustering displayed high sensitivity to noise, collapsing well below ARI = 0.3 and NMI = 0.3. HDBSCAN was more robust to mild Gaussian noise (**Figure 2A, 2B, 2C**) than other methods, except when compared to TopoTome.

Across increasing Gaussian noise levels (σ = 0.01-0.1), TopoTome maintained the mean ARI ≈ 0.95 and NMI ≈ 0.80. All other algorithms, including HDBSCAN, deteriorated substantially with increasing noise level, plummeting after σ = 0.04. Moreover, compared to TopoTome, all methods exhibited strong instability depending on the type of noise distribution (**Figure 2C, 2D, Supplementary figure 2B**). Runtime analysis showed that while TopoTome matched spectral and agglomerative clustering on small datasets (10^4^ voxels), but it scaled less efficiently at larger volumes (10^6^ voxels) (**Supplementary figure 3**). We could attribute this deficiency to the Leiden clustering step, which consists of multiple iterations of community detection over the whole 3D volume.

**Figure 3.**
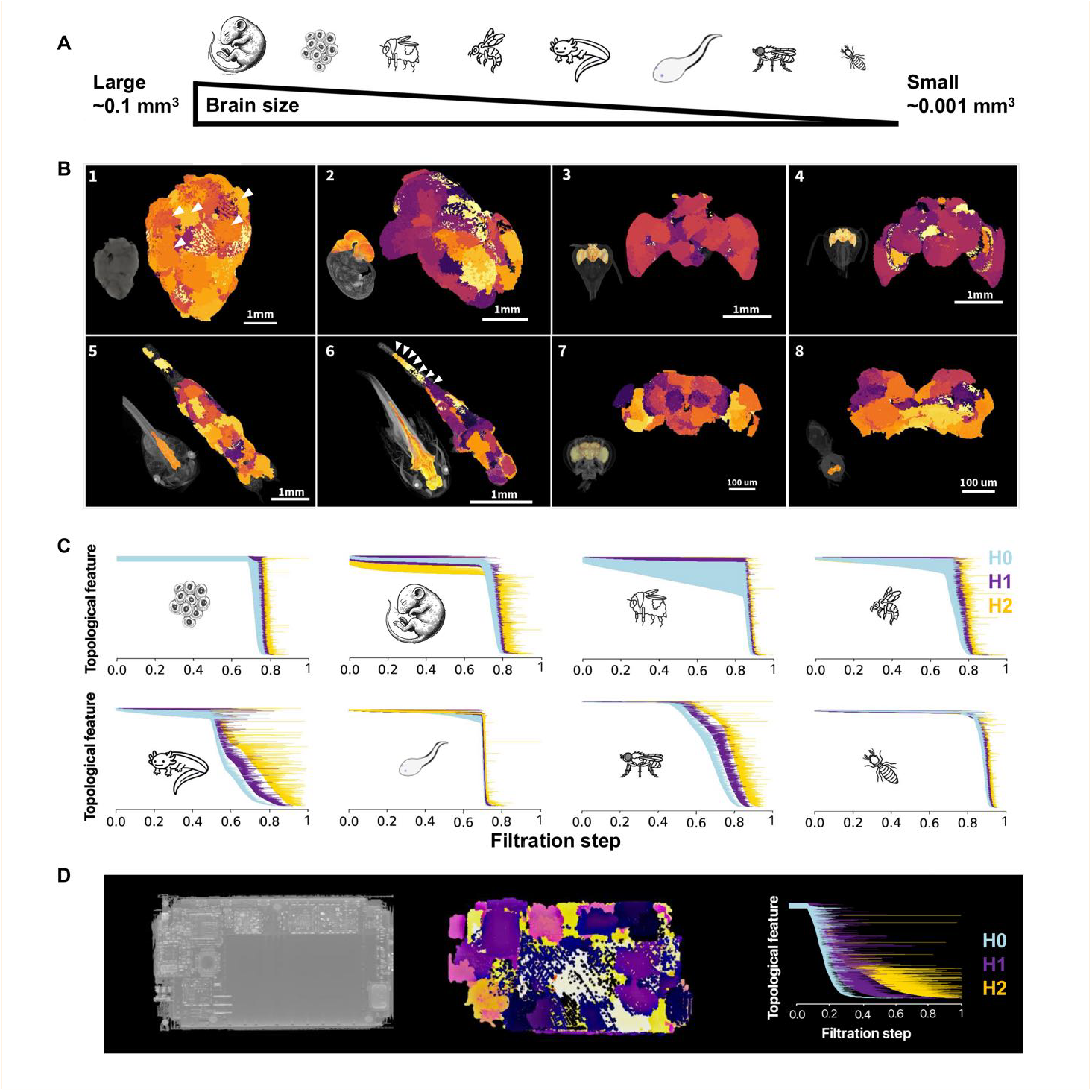
TopoTome generalizes across biological and non-biological data. **A)** Analyzed species sorted by their brain/organoid volume. **B)** TopoTome persistence cluster segmentation as applied to *μ* -CT scans of brains of various shapes and sizes. Panels 1: human brain organoid, 2: *M. musculus* E13.5 embryo, 3: *A. mellifera*, 4: *B. terrestris*, 5: *X. laevis* tadpole, 6: *A. mexicanum* tadpole, 7: *D. melanogaster*, 8: *R. speratus*. Lower left insets in panels: projection of the entire scanned volume with highlighted neuronal tissue that was segmented. White arrows point to organoid ventricles (panel 1) and vertebral column segments (panel 6). **C)** Persistence barcodes of brain structure from B (matching panel order). **D)** *μ*-CT scan, TopoTome segmentation, and persistence barcode of a smartphone (left to right).

These results demonstrate that topological analysis preserves meaningful structural features even when the signal is degraded. This is further supported by the performance of ToMATo, which also uses topological data analysis for image segmentation. ToMATo performed robustly under speckle and Poisson noise, reaching ARI and NMI values close to 0.8 - 0.9. However, unlike TopoTome, it exhibited reduced accuracy with Gaussian and Salt & Pepper perturbations (typically below 0.5), which are the most common challenges in bio-image data (**Figure 2D, Supplementary figure 2**). Crucially, we think that the better performance of TopoTome can be attributed to its implementation of the direct topological encoding and analysis of the entire voxel set, instead of transforming the data to subsampled point clouds as is implemented in ToMATo.

### Generalizability of TopoTome segmentation across biologically diverse subjects

We initially developed TopoTome to address the challenge of comparing morphologically diverse Drosophila brain regions imaged using X-ray micro-Computed Tomography. This task was challenging due to the absence of clearly comparable anatomical landmarks that could be used for manual segmentation and training a DL segmentation model (*μ*-CT; Bourgeat et al. *unpublished*). With TopoTome, we were able to delineate and compare fly brain tissue domains that broadly corresponded to the classical neuropil structures. In addition, TopoTome tiled the individual fly brains with finer, unique substructures that were not apparent to the naked eye, but were functionally relevant (**Figure 1B, Figure 3B**, panel 7; Bourgeat et al. *unpublished*). We next asked whether this approach could generalise beyond Drosophila. We found that the identification of brain regions with TopoTome can be naturally applied to any *μ*-CT brain tomography of diverse species with far more complex, larger, and different brain structures. This included brains from eusocial insect lineages, developing amphibians and mammals, and even human brain organoids **(Figure 3A)**. Besides brain structure, TopoTome could identify clusters corresponding to vertebral segments of the axolotl spinal chord (**Figure 3B**, panel 6) and organoid ventricles, well-known organoid and tissue structures that were not discernible to the human eye in the original image (**Figure 3B**, panel 1). The persistence barcodes of these brain structures revealed the diversity of brain topology across the animal kingdom, which can be further interrogated using topological data analysis (**Figure 3C**). Beyond biological systems, and without any modification of the algorithm, TopoTome was also able to segment non-biological 3D structures, such as the electronic components of a smartphone (**Figure 3D**). This was possible because TopoTome analyzes how voxels are arranged and connected locally, and in the context of the entire image, rather than relying on the image’s visual content. This makes it naturally generalizable to any image modality or subject.

While these segmentations appeared plausible, due to the low image resolution, human-annotated ground truths were not reliable or available for validation, while synthetic datasets may be overly simplified. To rigorously test the reliability of TopoTome segmentation, we therefore turned to published real-world 3D image benchmark datasets with established human-derived ground truth segmentations^7^.

### Benchmark against deep learning segmentation approaches

Unsupervised clustering methods are known to fare well with simple datasets but struggle with real-life images. In such cases, the standard approach has been to use a deep learning image segmentation model, trained on the manual annotations of a smaller subset of the data (usually subsets of 2D slices from a 3D image). We benchmarked TopoTome against such commonly used DL algorithms for 3D image segmentation using a more challenging, real-life 3D dataset^7^. The performance of these algorithms typically depends strongly on the data type on which they were trained. TopoTome is agnostic of the image data content or imaging modality, so it should be ubiquitously natively performant and applicable for segmentation of any dataset. We therefore compared TopoTome’s segmentation performance on the same datasets used to train the deep learning models we benchmark against. The data consisted of fluorescence microscopy images obtained at cellular resolution. Notably, this was not the 3D image data modality that TopoTome had been initially developed for. We also did not modify the TopoTome algorithm to optimize it for this dataset in any way.

We tested TopoTome segmentation on the standard 3D mouse skull nuclei dataset^7^ (**Figure 4A**). TopoTome achieved competitive F1 scores in the intermediate IoU range (τ = 0.3 - 0.5), where biological ground truth is most informative, with mean F1 values overlapping those of the top supervised pipelines (**Figure 4B**). Specifically, TopoTome performed within the range of the best-performing models, such as WNet3D (zero-shot) and StarDist with post-processing. Across some thresholds, it also outperformed CellPose and CellSeg3D (**Figure 4B, Figure 4D**). Unlike the deep learning methods that were specifically optimized for this type of dataset, TopoTome was applied out of the box. Its performance did not rely on training data, parameter optimisation, or post-processing, yet it stably outcompeted most other methods. TopoTome underperformed on runtime compared to the DL methods. However, executing TopoTome vs. a deep learning model is conceptually different (**Figure 1A**). Time efficiency of DL models is usually only assessed for the segmentation step, after ground truth generation and training for the DL models. Yet these multiple, time-consuming manual steps do not exist in TopoTome workflow. Hence, it becomes difficult to fairly compare the two in terms of time efficiency. We argue that the timeframe of TopoTome’s computation can be offset, at least in part, by the complete absence of ground truth annotation, model training, and optimization. Nevertheless, its computational footprint will need to be optimized.

**Figure 4.**
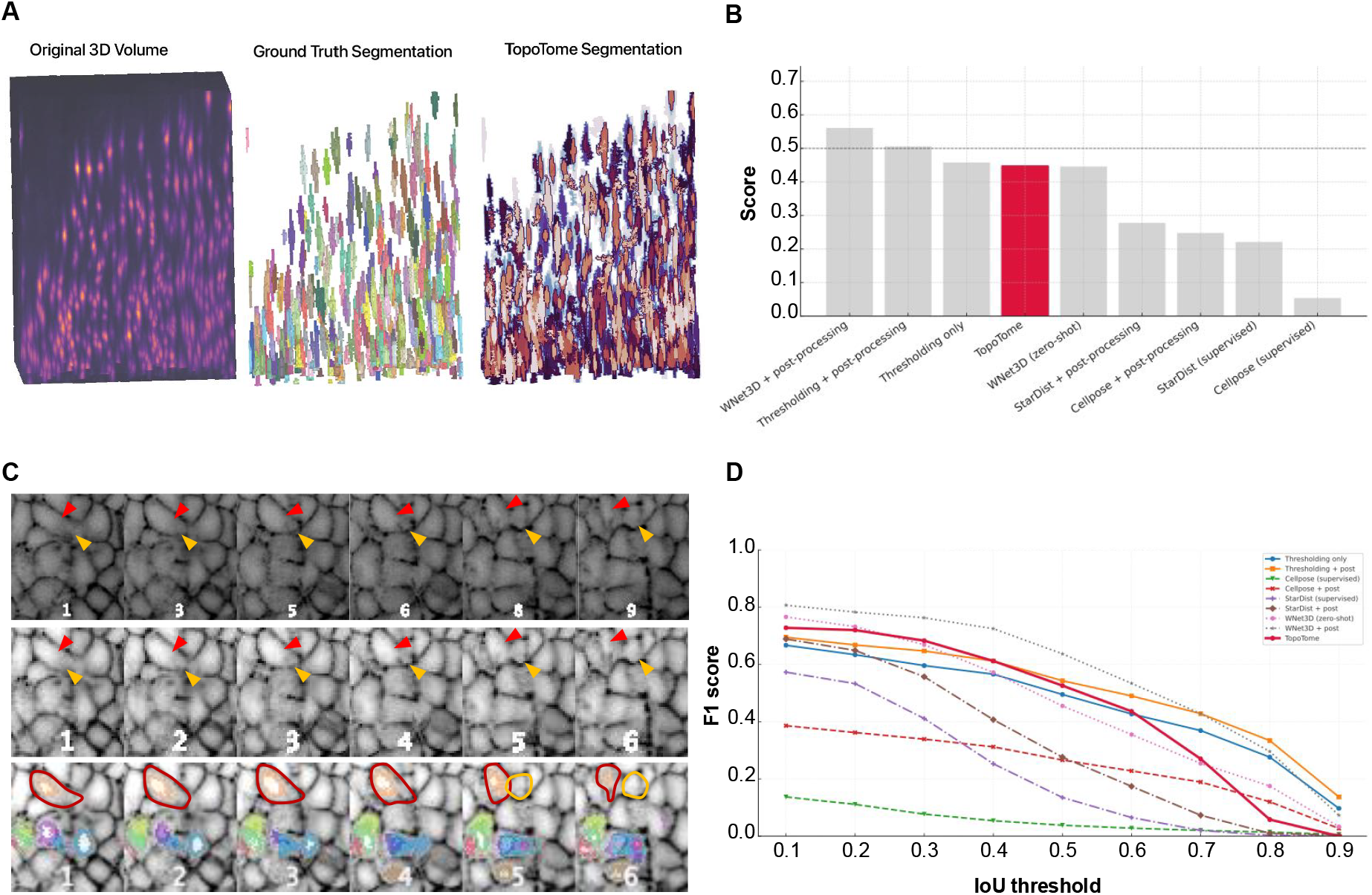
TopoTome segmentation generalizes across imaging modalities and complex biological data without training, tuning, or priors. **A)** Clustering of mouse neuron soma imaged using 3D confocal microscopy. Left panel: the original 3D image. Middle panel: the human-annotated ground truth image. Right panel: TopoTome segmented cluster outputs. **B)** Benchmarking TopoTome segmentation performance (F1 score) on an image from A) against tuned thresholding methods and DL models trained on equivalent datasets. TopoTome is applied “out-of-the-box” with no tuning or manual post-processing. **C)** An example of TopoTome segmentation of dense zebrafish embryo cells with low-resolution images depicts how TopoTome correctly segments a cell where human annotation fails. Top panels are a numbered sequence of 2D slices. The middle panel shows the equivalent sequence of 2D slices, but after isotropically downscaling the resolution to 60%. In the lower panel, a subset of TopoTome segments is shown in different colours, overlaid over the downscaled image data. Red and yellow circles depict human-annotated segmentation. Arrows point to the two true cells corresponding to what would be the correct annotation. **D)** Mean F1 score across IoU thresholds. TopoTome outperforms DL models across most IoU thresholds without relying on manual pre- or post-processing.

Although TopoTome is not yet optimized for efficient computation that would scale to very large image data, due to its conceptually different engine, it is accurate, generalizable, robust to noise, and autonomous. This makes it well-suited to generate mathematically defined ground truth for complex biological structures. We propose that this ground truth could be used to train DL models for a more accurate and human-independent segmentation workflow (blue pointers in **Figure 1A**). To this point, it is worth noting that after using TopoTome, we have observed in several cases across datasets that the human ground truth may not be correct. This is illustrated best in an example of TopoTome segmentation applied out-of-the-box to yet another type of dataset, a light-sheet microscopy of zebrafish embryo (**Figure 4C**). Especially at lower-resolution, high-content images such as densely packed zebrafish embryo cells, between-cell boundaries often escape detection when viewed one 2D slice at a time. This is, nonetheless, a standard way to manually segment 3D images. In our example in Figure 4C, only when the full volume context is taken into account, it may become apparent that the initially single segmented cell is, in fact, two cells with blurred boundaries. Such observations are anecdotal - we have not quantified this systematically because these kinds of errors are inherently not trivially noticed. To this point, we find that the F1 score tends to drop with increasing IoU threshold for both TopoTome and DL models. This means that the segmented objects match the ground truth, but their precise boundaries do not. Curiously, though, compared to DL models, this drop is distinctly sharper for TopoTome after *IoU >* 0.6 (**Figure 4D**). Because DL models are trained and dependent on ground truth, their F1 score decline across IoU thresholds is expected to be gradual, and proportional to the uncertainty of defining an accurate ground truth. This is not the case for TopoTome, where the definition of segment boundaries is deterministic. We therefore believe that this drop can be, at least partially, explained by variability and errors in human-defined ground truth, as seen in the example of the zebrafish embryo. This hypothesis is made more plausible by the fact that TopoTome outperforms DL models at lower thresholds.

Because of this, we envision that combining topological segmentation as a ground truth generator with DL and AI automation could be used for fully automated, scalable, and accurate bioimage analysis, even for the most challenging datasets, such as high-resolution brain imaging.

## Discussion

Our study highlights TopoTome, a topological encoding of 3D images, and a fully data-driven framework for quantitative comparative analysis of 3D images. Thanks to its topological data analysis core, it is robust to various sources of noise and stably performant, despite being agnostic to the imaged system. Besides 3D segmentation, TopoTome also provides topological encoding and quantification of the complete image, as well as local topological analysis of the segmented substructures. These outputs enable in-depth quantitative structural analysis of salient features beyond conventional volumetric measurements.

The main current limitation of TopoTome is its computational footprint and runtime, which is significantly slower than for pre-trained state-of-the-art DL-based segmentation algorithms. However, when factoring in the hundreds of hours typically required to prepare and validate training datasets for deep learning, TopoTome may prove to be comparatively more time-efficient than reported in this study. Unlike deep-learning-based segmentation models, TopoTome is not constrained by the subjectivity of human-annotated training data. Its segmentation is directly derived from the data’s mathematically defined structural properties.

We think the deficiencies of both TopoTome and deep-learning approaches are, in fact, complementary and can be overcome by playing into their respective strengths. For example, TopoTome can generate deterministic, mathematically defined ground truths to train fast and computationally efficient DL segmentation models (Figure 1A). Similarly, topological signatures of imaging artefacts could in principle be used for enhanced design of de-noising DL algorithms. Such synergy could transform the automation of designing and training neural networks used for bio-image analysis.

While TopoTome is still undergoing optimization of its computational footprint, it already offers a conceptually distinct approach to image data encoding, analysis, and segmentation, firmly grounded in topological data analysis. Because of this, it has the potential to address challenges that have been difficult to solve by tuning or re-imagining existing DL segmentation architectures.

TopoTome’s large computational footprint is, at its core, an engineering and not a conceptual problem. The current Python implementation, especially Leiden clustering, uses memory and processing resources sub-optimally. Future implementations in lower-level programming languages with improved memory management and parallelisation could substantially decrease the runtime and raise image size limits (currently capped to only a few GB tif stacks). Additionally, planned adjustments to the underlying TDA computation should remove the reliance on Leiden clustering, which is set to significantly boost its efficiency in the future.

While we emphasized the segmentation utility of TopoTome as its most immediate application, its broader strength lies in encoding entire structural volumes from raw image data directly into the topological space. By allowing topological data analysis on the complete image topology, TopoTome shifts bio-structure analysis from relative comparisons of meshes, volumes, or derived networks toward topological quantification of the complete shape of virtually any 3D structure. Topological data analysis, which is at the heart of the TopoTome algorithm, has no theoretical limitations on encoding or analysing structural complexity. This capability holds promise for advancing our understanding of structurally most intricate biological systems.

The first version of TopoTome provides a conceptually fundamental shift in structural image analysis. With future improvements to its implementation and efficiency in handling large images, we can envisage we could use it to understand even the most complex structures, including the complete ultrastructure of the brain.

## Materials and Methods

### Topological data analysis underlying TopoTome

TopoTome implements an algorithm that uses the persistence diagram to extract local geometrical information from global topological space in the form of local persistence homologies. Here, we propose an algorithmic approximation of the calculation of local homologies. Our approach aims at finding a mapping *ϕ* that takes a pair of persistent points from the persistence diagram, defined by birth *b* and death *d* values of the topological feature (as described in *Cubical complex filtration*), and maps them to the associated cluster of voxels in the original image (ℤ^3^):

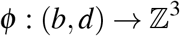

Let *I*: ℤ^*n*^ → ℝ represent a greyscale image, where ℤ^*n*^ denotes the image’s voxel grid in *n* dimensions (typically *n* = 3 for the 3D images), and ℝ represents the intensity value at each voxel. A *cubical complex 𝒦* (*I*) is a topological space constructed from the image by associating a cube (or hypercube in higher dimensions) with each voxel. Formally, the cubical complex *𝒦* (*I*) is defined as the union of all elementary cubes *C*_*i*_ associated with the voxels in the image:

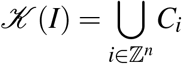

Each *C*_*i*_ corresponds to a unit cube centred at the voxel *i* with a dimension corresponding to the number of non-zero entries in the binary representation of *i*. We perform a specific filtration over the data (cubical complex filtration) to eventually identify the persistent features in the image. Formally, a *filtration* is a nested sequence of subcomplexes {*𝒦*_*α*_ (*I*) _*α∈*_ ℝ such that *𝒦*_*α*_ (*I*) ⊆ *𝒦* (*I*) for *α* ≤ *β*. For a cubical complex derived from a grayscale 3D image *I*, the filtration is induced by the voxel grayscale intensity values. We, hence, define the sub-complex *𝒦*_*α*_ (*I*) as:

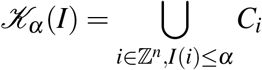

Here, *𝒦*_*α*_(*I*) contains only the cubes *C*_*i*_ whose corresponding voxel intensity *I*(*i*) is less than or equal to *a*. We then compute the k-th homology group of each subcomplex over each filtration step (intensity threshold step or “time” steps) as follows. Let *H*_*k*_(*𝒦*_*α*_ (*I*)) denote the *k*-th homology group of the subcomplex *𝒦*_*α*_ (*I*). The rank of this group, *β*_*k*_(*𝒦*_*α*_(*I*)), is called the *k*-th Betti number and counts the number of *k*-dimensional features (connected components, loops, voids). The *persistent homology* tracks the changes in these homology groups as *a* varies. To define the persistence diagram, we first define *p*_*k*_(b, d) as a persistent *k*-dimensional feature, where b is the birth time and d is the death time. The collection of such features forms the *persistence diagram D*(*I*) of the image *I*:

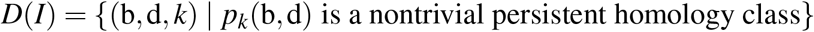

We subset the persistence diagram, define the function that takes a persistent feature, and map it onto the image. Let *D*_2_(*I*) denote the set of 2-dimensional persistence points in the persistence diagram *D*(*I*). Each point (*b, d*) ∈ *D*_2_(*I*) represents a 2-dimensional topological feature (a void) with birth time *b* and death time *d*:

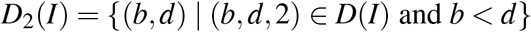

For each persistence point (*b, d*) ∈ *D*_2_(*I*), we associate a specific cluster of voxels within the image and define the cluster of voxels associated with a persistence point (*b, d*) as a set *V*_(*b,d*)_ ⊆ ℤ^*n*^. Finally, we define the mapping 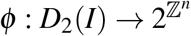 that assigns to each 2-dimensional persistence point (*b, d*) a corresponding cluster of voxels *V*_(*b,d*)_:

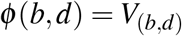

Here, 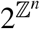 denotes the power set of ℤ^*n*^, which is the set of all possible voxel clusters. Since the number of persistence points and voxel pairs within any persistent interval (*b, d*) is |*V*_(*b,d*)_| ≥ 1 for any pair of persistence points, we combined our approach with the application of the Leiden graph clustering algorithm^28^. More precisely, to characterise the voxel cluster *V*_(*b,d*)_, we apply the Leiden algorithm to the set of voxels *P*_(*b,d*)_ ⊆ ℤ^*n*^ involved in the 2-dimensional feature associated with (*b, d*) after transforming the voxels into a point cloud dataset. The Leiden algorithm is then applied to *P*_(*b,d*)_ to identify clusters of voxels:

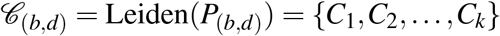

From the obtained partition 𝒞_(*b,d*)_, we select the most significant community and define it as the actual cluster. Hence, we define a function *Ψ*: 𝒞 _(*b,d*)_ → {ℝ} that evaluates each cluster *C*_*i*_ for community size and select the conform cluster *V*_(*b,d*)_ as:

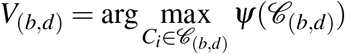

The final mapping of a 2-dimensional persistence point (*b, d*) to a voxel cluster *V*_(*b,d*)_ in the image is given by:

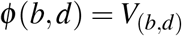

where *V*_(*b,d*)_ is the cluster of voxels obtained by applying the Leiden algorithm to *P*_(*b,d*)_ and selecting the largest cluster. Each such defined cluster of voxels is enumerated and considered a distinct substructure of the analyzed 3D volume (“*persistent cluster*”; see Figure 1B and D). The minimal absolute volume of the substructure should be limited depending on the voxel resolution. The default relative size is arbitrarily limited to 50 voxels (initially used to capture anatomically significant topologically distinct structures in a fly brain micro-CT image).

### Calculation of local topology of persistent clusters

Persistent clusters are often numerous, so their structure often cannot be represented efficiently by calculating as many persistent barcodes. Therefore, after extracting the persistence clusters, we approximate the clusters’ homeomorphism to the *n*th-homology groups using the Euler Characteristic Transform^29–31^. The Euler Characteristic is defined as the following alternating sum:

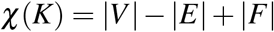

where *K* is a polyhedron and |*V*|,|*E*|, |*F*| are the number of vertices, edges and faces, respectively. It is important to note that the Euler characteristic does not provide the complete homological information on the structure as the persistence diagram does, but unlike the persistence diagram approach, ECC exhibits linear instead of exponential computational complexity, making it more suitable for local homology approximation for many sub-volumes^29–31^. For a simplicial complex of a polyhedron *K*, the Euler characteristic 𝒳 (*K*) is defined as:

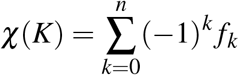

where:

- *f*_*k*_ is the number of *k*-dimensional simplices in *K*,
- *n* is the dimension of the highest-dimensional simplex in *K*.

In terms of homology, the Euler characteristic can also be expressed using the Betti numbers *β*_*k*_, which represent the rank of the *k*-th homology group *H*_*k*_(*K*). The Euler characteristic is then given by:

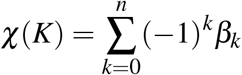

where *β*_*k*_ counts the number of *k*-dimensional holes in the complex. In image analysis with simplicial or cubical complex filtration, we have different subcomplexes at each filtration point. To follow the evolution of the Euler Characteristic across the filtration, we define the Euler characteristic curve (ECC) as the function:

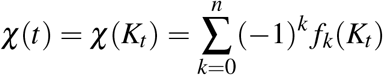

where *t* is the filtration step. This curve approximates the homology of the simplicial or cubical complex by summarising its topological features. The Euler characteristic only captures the *alternating sum* of the Betti numbers. Via the Gauss-Bonnet theorem, we know that the Euler characteristic is proportional to the overall Gaussian curvature of the studied shape, making it an advantageous morphometrics approach. From the local Euler Characteristic curves we can define the *Euler characteristic transform* (ECT) for each cluster, denoted by 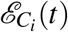, as:

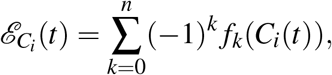

where *C*_*i*_(*t*) is the filtered subcomplex associated with the cluster *C*_*i*_ at filtration value *t*, and *f*_*k*_(*C*_*i*_(*t*)) represents the number of *k*-dimensional simplices in the cubical complex of the cluster. This procedure can also be used to analyse point cloud datasets represented by coordinate matrices of the persistent cluster, with the resulting topological spaces being referred to as simplicial complexes^32,33^. This allows one to measure the shape of the cluster, rather than its internal structure. TopoTome outputs the cubical-complex derived Euler Characteristic Curve for every detected segment as a measure of their local structure.

### Algorithm implementation

TopoTome is implemented in the Python programming language. Full implementation, installation, and usage guidelines, including a dependency environment, and Jupyter notebooks with examples, can be found on the TopoTome git repository. The main pipeline used in this paper is implemented in the *runner*.*py* script, which executes image import and processing, cubical complex filtration of the volume, persistent homology lifespan calculation, identification of persistence clusters, and their mapping to the original voxels. A Python script *homology2voxel*.*py* implements the main analytical part of the algorithm. It uses the *CubicalRipser* Python package as the backbone for topological data analysis, specifically for the cubical complex filtration^34^ and the *leidenalg* Python package for Leiden clustering,^28^. The standard input file for TopoTome is a stack of grayscale 2D images in.tiff format. Output files consist of a.pkl file containing the persistence diagrams and a folder containing.h5 files for every identified persistence cluster, i.e. segmented object. The.h5 file contains a binary mask of the cluster as z, x, y coordinates that correspond to voxel coordinates of the original input TIFF image. The files containing segmented cluster masks can be visualised with respect to the original 3D volume with the *visualisation*.*py* script. Euler Characteristic Curves corresponding to each persistent cluster (segment) are output in a.csv table.

### Synthetic datasets

We first generated synthetic data using a set of random 2D Gaussian blobs with varying voxel intensity to mimic real biological *μ*-CT images. We then added Gaussian noise (*ε* ∼ 𝒩 (*μ, σ*), where *μ* is generally 0) to the whole volume.

We synthesised five 3D images composed of 3D Gaussian blobs per increasing noise level, ranging from *σ* = 0.01 to *σ* = 0.1.

### Clustering performance benchmark

For benchmarking, we compared TopoTome’s performance with established clustering methods: K-Means, HDBSCAN, agglomerative clustering, spectral clustering, and ToMATo (a topological data analysis-based point-cloud clustering algorithm^19^). These methods are implemented within the Scikit-learn and tomaster libraries.

Since TopoTome operates natively on image volumes, while the other algorithms require point cloud input, we constructed feature vectors from the synthetic images. The synthetic volumetric images were converted into point clouds by applying an intensity threshold of 0.1, retaining only voxels above this value. This thresholding step reduces background noise while preserving meaningful structures, thereby ensuring consistent and comparable input representations across all clustering methods.

For 2D datasets, feature vectors were built solely from voxel coordinates. For 3D datasets, we prepared two types of feature vectors: one based only on coordinates, and another combining coordinates with voxel intensity values, in order to account for algorithms that exploit intensity information.

Each clustering method (K-Means, HDBSCAN, agglomerative clustering, spectral clustering, and ToMATo) was applied to the same synthetic datasets, with clustering performance evaluated against ground truth labels. Robustness analyses were performed by varying noise levels (*s* between 0.01 and 0.1) and by repeating experiments across 100 independent runs for each algorithm. Computational complexity was assessed by running all algorithms on volumetric images of increasing size, ranging from 10^4^ to 10^6^ voxels.

To compare the performance across different segmentation software, we employed two widely-used metrics: the Adjusted Rand Index (ARI) and the Normalised Mutual Information (NMI). The ARI is a pair-counting measure that evaluates the similarity between two clusterings by considering all pairs of samples and correcting for chance agreement. It is defined as

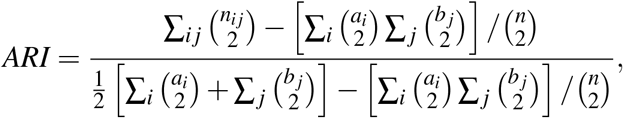

where *n*_*ij*_ is the number of samples shared by the *i*th true cluster and the *j*th predicted cluster, *a*_*i*_ = Σ _*j*_ *n*_*ij*_ and *b*_*j*_ = Σ_*i*_ *n*_*ij*_ are the cluster marginals, and 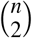 is the total number of sample pairs. NMI is an information-theoretic metric that quantifies the amount of information shared between the true clustering *U* and the predicted clustering *V*, normalised by the average of their entropies:

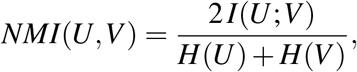

with *I*(*U*;*V*) denoting the mutual information between *U* and *V*, and *H*(*U*) and *H*(*V*) being the respective entropies. These metrics provide robust, label-invariant evaluations that capture both the local pairwise agreements (ARI) and the global similarity in cluster structure (NMI) between an algorithm’s output and the ground truth.

### Noise robustness and benchmarking against deep learning methods

To evaluate robustness under realistic imaging conditions, we introduced four types of noise into synthetic volumetric datasets: Gaussian noise (to model acquisition blur, *σ* = 0.01), salt & pepper noise (to emulate sensor dropout, parameter *θ* = 0.05), speckle noise (to mimic multiplicative distortions, *s* = 0.01), and Poisson noise (to reflect photon-limited environments, with default `scikit-image` parameters. These perturbations were selected to represent common degradation sources in microscopy and medical imaging pipelines^27^. All clustering algorithms, including TopoTome, were applied to these noise-augmented datasets under identical conditions.

In addition, we benchmarked TopoTome against supervised Deep Learning pipelines using the *Mouse-Skull-Nuclei-CBG* dataset^7^. This dataset consists of light-sheet microscopy images of nuclei in the mouse skull, with ground-truth annotations that provide instance-level labels for hundreds of nuclei. Competing models included segmentation architectures WNet3D and StarDist, with and without manual post-processing^7^. TopoTome was applied directly to the raw volumetric images without post-processing. To evaluate the performances of different deep learning methods and TopoTome, we used a unified evaluation framework. Namely, we used Intersection-over-Union (IoU)-based F1 scoring scheme, while systematically computing overlap between predicted clusters and ground-truth annotations across varying thresholds.

### Animal husbandry and ethics

A 5-day-old adult vinegar fly (*Drosophila melanogaster*) of a wild-type genotype from the Drosophila Genetic Reference Panel^35^ was selected from a batch of flies that was reared for 2 generations in a controlled environment and population density at 26^*o*^*C*, with a humidity of 50-70% and a day/night cycle of 12 h. The fly was decapitated, and the head was prepared for the micro-CT scan. The Japanese termite (*Reticulitermes speratus*) was reared in a standard laboratory colony on wood bark before getting anaesthetised and decapitated for scans. A single female honey bee (*Apis mellifera*) and a single buff-tailed bumblebee (*Bombus terrestris*) were both caught foraging on cherry laurel (Renens, Switzerland), on the morning of 12th June 2024. They were fed raspberry sugar syrup, then anaesthetised and decapitated for scans on the same day. Adult zebrafish (*Danio rerio*) were maintained under standard conditions on a 14 h/10 h light cycle at the accredited EPFL fish facility (VD-H23). *TgBAC(cdh2:cdh2-GFP)\**^**?**^ Zebrafish embryos were obtained by natural spawning. They were incubated at 28.5^*o*^*C* and shifted to 19.5^*o*^*C* after shield stage. The 15-somite stage embryos were imaged the next day.

Rearing of a juvenile (*Ambystoma mexicanum*) and all experimental procedures were performed according to the Animal Ethics Committee of the State of Saxony, Germany. All tadpole (*Xenopus laevis*) embryo samples were collected following the Swiss Federal Veterinary Office guidelines and as authorised by the Cantonal Veterinary Office (cantonal animal license no: VD3652c, and national animal license no: 33237). The mouse embryo (genotype WT CD1, stage 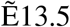) was collected in accordance with the Swiss Federal Veterinary Office guidelines and as authorised by the Cantonal Veterinary Office (cantonal animal license no: VD3652c, national animal licence no: 33237). Zebrafish husbandry procedures have been approved and accredited by the federal food safety and veterinary office of the canton of Vaud (VD-H23). No sex determination was performed in bumblebee, mouse, axolotl, zebrafish embryo, or tadpoles used in this study. Invertebrates used in this study were not subject to formal ethical approvals, but were nonetheless reared and handled with care.

### *μ*-CT and light-sheet microscopy sample preparation

To prepare the samples for CT scans, we used potassium iodide (*I*_2_*KI*, Lugol’s solution) to provide contrast staining in all tissues^36,37^). *I*_2_*KI* binds to alpha-1,4 glucans found ubiquitously in polysaccharides^37,38^, as well as to lipids, both being abundant inside the cell lumen and their membrane^39,40^. Heads or whole bodies of animals were placed in 1.5 mL Eppendorf tubes containing 1 mL of 0.5% Triton X-100 in 1x Phosphate Buffered Saline (0.5% PBST) for 5 min. PBST solution was replaced with Bouin’s solution for 24 hours for fixation at room temperature (21^*o*^*C* + −1^*o*^*C*). After the fixation, the sample was washed three times for 30 min in a µ-CT wash buffer (0.1 M Na2HPO4, 1.8% Sucrose) and incubated in 1 mL of *I*_2_*KI* (Lugol’s solution) at room temperature (21^*o*^*C* + −1^*o*^*C*) for smaller samples such as vinegar fly heads and termites. A 5-day incubation was used for the fly and termite samples, while a 10-day incubation was used for the honey bee, bumblebee, organoid, axolotl, frog tadpole, and mouse embryo samples. For even larger samples than those, empirical adjustment of the length of incubation is advised to allow for iodine to reach all internal structures of the sample of interest. Lugol’s solution was removed and samples were washed three times in ultrapure water, each for 30 min, following the recommendations from the literature^41^. We recommend skipping these three washing steps, as they reduce the bleed-out of the staining agent and increase imaging contrast. We mounted the samples in a p200 or p1000 pipette tip (depending on the size of the sample) filled with ultrapure water. We sealed the tips using dental wax (Thermo Fisher) to avoid water evaporation during the scans.

The zebrafish embryos were dechorionated and laterally placed on 2% low-melting-point agarose with depressions. Imaging was performed in fish water at 28.5 °C on a Viventis LS1 light-sheet microscope with a CFI75 Apochromat 25×, NA 1.1 water-immersion objective (Nikon). The light sheet was generated by scanning of a 2.2 *µm* diameter Gaussian beam at 488 nm, yielding an anisotropic voxel size (*x* = 0.347 *µm, y* = 0.347 *µm* and *z* = 0.5 *µm*). Detailed method can be found in^**?**^. The greyscale colours of the obtained image were inverted in ImageJ such that the stained membranes have low intensity. We then downsized it to a stack of 41 images (slice 100 to 140), downscaled to 80% or 60% size using voxel averaging without interpolation. We analyzed a rectangle selection of an arbitrary size (190 × 98 pixels, at 60% downscale) which was chosen also arbitrarily from around the centre of the image, only making sure it overlapped a region with densely packed cells.

### µ-CT scan parameters

The X-Ray µ-CT machine (Scanco Medical, µCT 100 cabinet µ-CT scanner) parameters were set to: 45 kVp (maximum high voltage applied to the X-ray tube), intensity to 88 µA, and power to 4 *W*. Integration time was set to ≈350 *ms*. A 0.5 *mm* aluminium filter was used to decrease the X-ray beams’ scattering. The images were reconstructed from a 3D matrix of density scales of images obtained by imaging the rotating sample (360^*o*^). The resulting voxel size was 3.3 *µ*m, isotropic. Projection images were averaged over three frames. Nano-CT volumes were acquired on the RX Solutions UltraTom with a 160kV nano-focus tube at 795 *nm* isotropic voxel size. We used an intensity of 125 *µA*, a voltage of 40 *kV p*, and an integration time of ≈ 1000 *ms*.

## Data availability

All data produced for this study (*µ*-CT scans, light-sheet zebrafish embryo data, and synthetic simulated data) can be downloaded from the Zenodo repository or from https://github.com/jaksiclab/TopoTome/.

µ-CT scan of the smartphone was downloaded as open source from https://dragonfly.comet.tech/en/sample-datasets^42^.

## Ethics statement

All animals subject to ethics regulations used in this study were licensed to be used and handled according to ethics regulations, as described in Methods: Animal husbandry and ethics.

## Acknowledgements

We would like to thank EPFL’s PIXE facility for the support with nano-CT scans, Riddha Manna and Benjamin Gallusser for helpful discussions, Pierre Gönczy for kindly donating the termite samples, Tatiana Sandoval Guzman for kindly donating the axolotl samples. This study was funded by the ELISIR scholarship and EPFL School of Life Sciences.

## Author contributions statement

S.S.B. and A.M.J. conceptualised the study, designed the experiments, analyzed and interpreted the data. S.S.B. prepared tissue samples, optimised staining procedures, and analysed the data. S.S.B. and V.G. wrote the TopoTome software. F.D. optimised the X-ray tomography and performed micro-CT head scans. E.N.B. and F.Z. provided the brain organoid sample. C.A. provided the axolotl, frog and mouse embryo samples. A.C.O and F.N.A. provided zebrafish embryo data. P.M.R. and A.M.J. supervised the study. S.S.B., F.D., P.M.R. and A.M.J. wrote the manuscript.

