## Supplemental figures and table for "TopoTome: Topology-informed unsupervised segmentation and analysis of 3D images"

### Supplementary figures

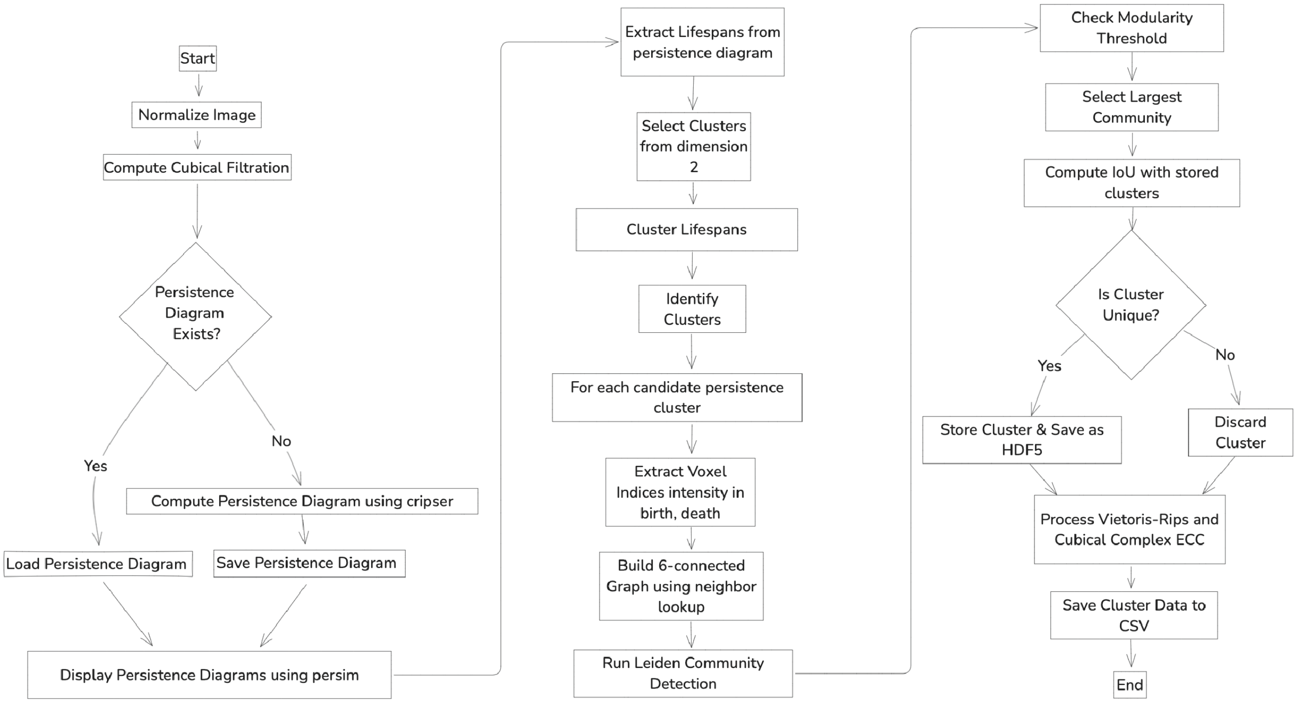


**Supplementary Figure 1. TopoTome algorithm.**

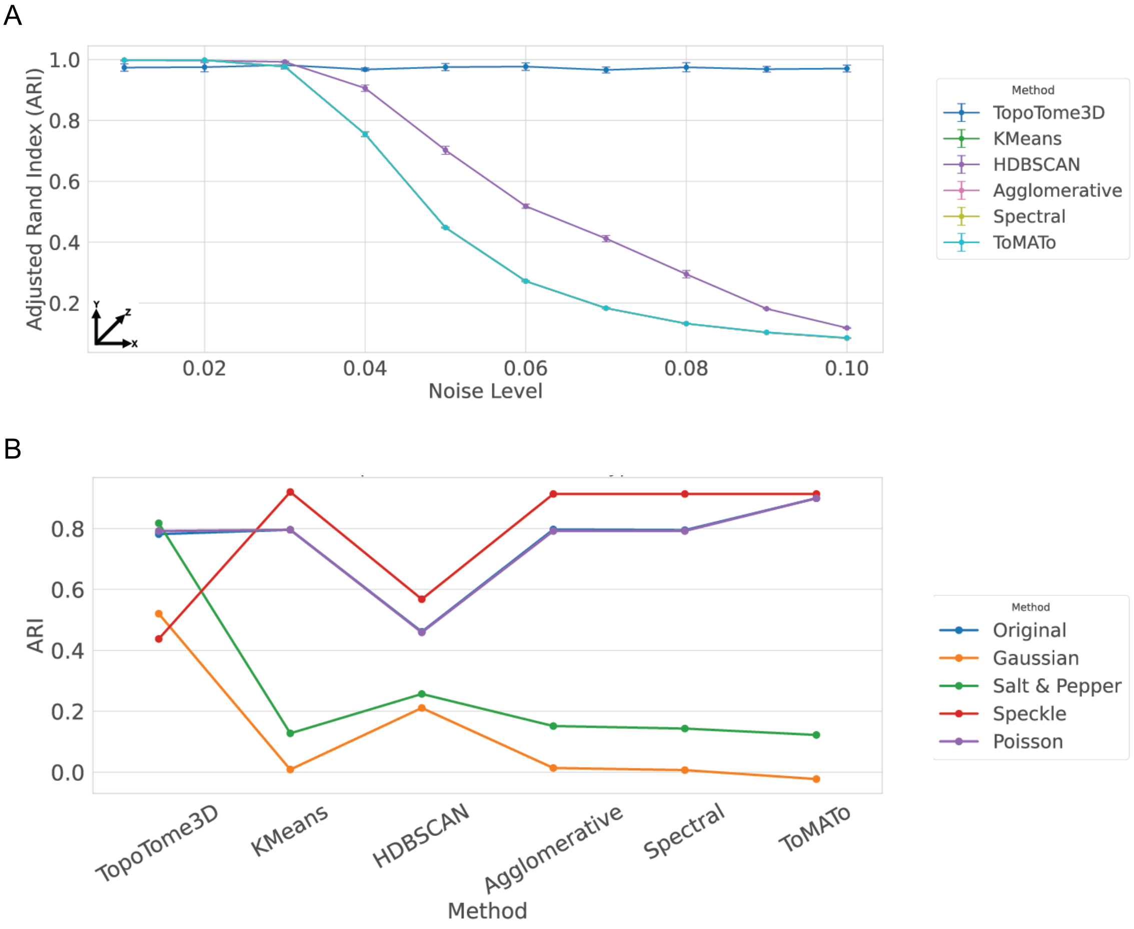


**Supplementary Figure 2. Adjusted rand indices across unsupervised clustering algorithms. A)**ARI across a range of Gaussian noise levels. **B)** ARI across different noise types.

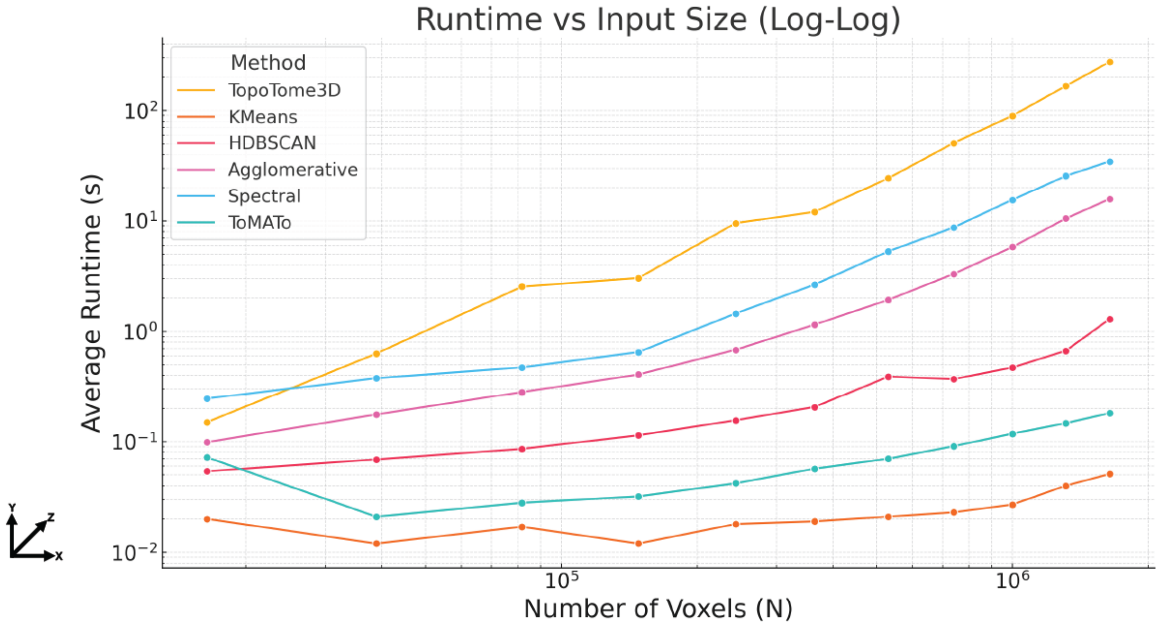


**Supplementary Figure 3. Runtime benchmark. A)** Average runtime in seconds across a range of voxels. Axes are shown in log scale.

### Supplementary tables

**Algorithm 1 TopoTome**

**1. Import and Normalize Image**
Load 3D image (TIFF)
Normalize intensities to range $[0,1]$

**2. Compute Persistence Diagrams from cubical complex filtration**
Compute persistence diagrams (PDs) using cubical complex filtration on the 3D greyscales
Extract persistence points' lifespans for each dimension

**3. Select Clusters Based on Lifespans**
Choose clusters from dimension 2
Compute lifespans: $\textit{lifespan} = \textit{death} - \textit{birth}$
Optionally select top $N$ clusters by largest lifespans

**4. Identify Clusters Using Leiden Algorithm**
**for** each selected cluster **do**
    Extract voxel indices where birth $\leq$ intensity $\leq$ death
    Build voxel graph (6-connectivity)
    Apply Leiden community detection on graph
    Select the largest community as the true cluster
    **if** size $\geq$ threshold and Intersection over Union (IoU) ensures uniqueness **then**
        Compute Euler Characteristic Curve (ECC)
        Store cluster data and ECC
    **end if**
**end for**

**5. Ensure Cluster Uniqueness**
Compute IoU for clusters
Retain unique clusters with IoU below threshold

**6. Store and Analyze Clusters**
Save cluster masks and ECC data for analysis
